# *Ehrlichia ruminantium* uses its transmembrane protein Ape to adhere to host bovine aortic endothelial cells

**DOI:** 10.1101/2021.06.15.447525

**Authors:** Valérie Pinarello, Elena Bencurova, Isabel Marcelino, Olivier Gros, Carinne Puech, Mangesh Bhide, Nathalie Vachiery, Damien F. Meyer

**Affiliations:** CIRAD, UMR ASTRE, F-97170 – Petit-Bourg, Guadeloupe, France; ASTRE, CIRAD, INRA, Univ Montpellier – Montpellier, France; Laboratory of biomedical microbiology and immunology, University of veterinary medicine and pharmacy in Kosice, Komenskeho 73 – Kosice, Slovakia; Institut Pasteur de la Guadeloupe – Les Abymes Cedex, Guadeloupe, France; Institut de Systématique, Évolution, Biodiversité (ISYEB), Muséum national d’Histoire naturelle, CNRS, Sorbonne Université, EPHE, Université des Antilles Campus de Fouillole – Pointe-à-Pitre, France; Institute of neuroimmunology, Slovak academy of sciences, Dubravska cesta 9 – Bratislava, Slovakia

**Keywords:** Intracellular bacterium, *Ehrlichia*, pathogenesis, adhesion, mammalian host cell, vaccine candidate

## Abstract

*Ehrlichia ruminantium* is an obligate intracellular bacterium, transmitted by ticks of the genus *Amblyomma* and responsible for heartwater, a disease of domestic and wild ruminants. High genetic diversity of *E. ruminantium* strains hampers the development of an effective vaccine against all strains present in the field. In order to develop strategies for the control of heartwater through both vaccine and alternative therapeutic approaches, it is important to first gain a better understanding of the early interaction of *E. ruminantium* and its host cell. Particularly, the mechanisms associated with bacterial adhesion remain to be elucidated. Herein, we studied the role of *E. ruminantium* membrane protein ERGA_CDS_01230 (UniProt Q5FFA9), a probable iron transporter, in the adhesion process to host bovine aortic endothelial cells (BAEC). The recombinant version of the protein ERGA_CDS_01230, successfully produced in the *Leishmania tarentolae* system, is O-glycosylated. Following *in vitro* culture of *E. ruminantium* in BAEC, the expression of CDS ERGA_CDS_01230 peaks at the extracellular infectious elementary body stages. This result suggest the likely involvement of ERGA_CDS_01230, named hereafter Ape for Adhesion protein of *Ehrlichia*, in the early interaction of *E. ruminantium* with its host cells. We showed using flow cytometry and scanning electron microscopy that beads coated with recombinant ERGA_CDS_01230 (rApe) adheres to BAEC. In addition, we also observed that rApe interacts with proteins of the cell lysate, membrane and organelle fractions. Additionally, enzymatic treatment degrading dermatan and chondroitin sulfates on the surface of BAEC is associated with a 50% reduction in the number of bacteria in the host cell after a developmental cycle, indicating that glycosaminoglycans seem to play a role in the adhesion of *E. ruminantium* to the host cell. Finally, Ape induces a humoral response in vaccinated animals. Globally, our work identifying the role of Ape in *E. ruminantium* adhesion to host cells makes it a gold vaccine candidate and represents a first step toward the understanding of the mechanisms of cell invasion by *E. ruminantium*.

## Introduction

*Ehrlichia ruminantium* is an obligate intracellular Gram-negative bacterium responsible for the fatal and neglected heartwater disease of domestic and wild ruminants (Allsopp, 2010). This bacterium belongs to the *Anaplasmataceae* family in the order Rickettsiales that includes many pathogens and symbionts of veterinary and public health importance (Moumene and Meyer, 2015). *E. ruminantium* is transmitted by ticks of the genus *Amblyomma* in the tropical and sub-Saharan areas, as well as in the Caribbean islands. It constitutes a major threat for the American livestock industries since a suitable tick vector is already present in the American mainland and potential introduction of infected *A. variegatum* through migratory birds or uncontrolled movement of animals from Caribbean could occur (Deem, 1998; Kasari et al., 2010). The disease is also a major obstacle to the introduction of animals from heartwater-free to heartwater-infected areas into sub-Saharan Africa and thus restrains the breeding to upgrade local stocks (Allsopp, 2010). The small genome of *E. ruminantium* (1.5 Mb) shows an unique process of contraction/expansion in non-coding regions and targeted at tandem repeats (Frutos et al., 2006). This genome plasticity, associated with a high genetic diversity, suggests a capacity of adaptation upon exposure to a novel environment and could also explain the low field efficacy of available vaccines (Cangi et al., 2016).

After adhesion and entry of infectious elementary bodies into host cells, *E. ruminantium* replicates by binary fission of reticulated bodies into an intracellular vacuole bounded by a lipid bilayer membrane derived from the eukaryotic host endothelial cell membrane (Dumler et al., 2001). *Ehrlichia* spp. have evolved sophisticated mechanisms to invade and multiply in host tissues by hijacking/subverting host cell processes ranging from host signaling, modulation of vesicular traffic, protection from oxidative burst, acquisition of nutrients, and control of innate immune activation (Moumene and Meyer, 2015). Notably, *E. chaffeensis* secretes the type IV effector Etf-1 to induce autophagy and capture nutrients, whereas it uses Etf-2 to delay endosome maturation to avoid phagolysosomal fusion for the benefit of bacterial replication (Lin et al., 2016; Yan et al., 2018). Moreover, recent work identified that *E. chaffeensis* uses EtpE invasin to enter mammalian cells via the binding to its receptor DNaseX, a glycosylphosphatidylinositol-anchored cell surface receptor (Mohan Kumar et al., 2013). That receptor-triggered entry simultaneously blocks the generation of reactive oxygen species (ROS) by host monocytes and macrophages (Teymournejad et al., 2017).

Due to the lack of some key metabolic genes that are required for host-free living and similarly to what is observed in other intracellular bacteria, entry into the eukaryotic host cells is crucial for *E. ruminantium* to sustain its life and disseminate (Pizarro-Cerda and Cossart, 2006; Frutos et al., 2007). Computational studies have predicted type IV effectors in *E. ruminantium* (Noroy et al., 2019) which, however, remain to be characterized; the mechanisms of adhesion and entry are still unknown and may be active or passive depending on the pathogen. *E. ruminantium* could either inject a type IV effector across the host cell membrane to trigger actin rearrangement and pathogen phagocytosis such as *Bartonella henselae* (Truttmann et al., 2011). An another option is the use of homologous system of *E. chaffeensis* invasion/receptor pair (Mohan Kumar et al., 2013) or an outer membrane protein of the OmpA family that actively trigger internalization as was observed in *Coxiella burnettii* (Martinez et al., 2014).

Bacterial pathogens are capable of exploiting and diverting host components such as proteoglycans for their pathogenesis (Aquino et al., 2018). These assemblies of glycosaminoglycans (GAGs) chains fixed around a protein nucleus (Bartlett and Park, 2010) are expressed constitutively on the cell surface, in intracellular compartments as well as in the extracellular matrix. They are known to act as receptors for pathogens in many cases of infection (Aquino et al., 2018; Rajas et al., 2017; Gagoski et al., 2015). Many pathogens – *e.g. Chlamydia trachomatis* with a biphasic developmental cycle like *E. ruminantium* – use GAGs as an initial anchor site of low affinity; this facilitates interaction with their respective secondary receptor allowing internalization but GAGs are sometimes also used as bridging molecules (Aquino et al., 2010). Thus, whether *E. ruminantium* uses GAGs as a portal of entry or any specific bacterial surface protein is unknown but essential for developing any anti-infective measures.

The outer membrane proteome study of *E. ruminantium* Gardel strain revealed that a hypothetical protein, the possible major ferric iron binding protein precursor the putative iron transporter ERGA_CDS_01230, is uniquely expressed in the outer-membrane fraction (Moumene et al., 2015). This protein was also shown to be O-glycosylated only in *E. ruminantium* (Marcelino et al. 2019). Moreover, homologous counterparts of this protein in other pathogenic species play a key role in bacterial survival within the host by scavenging iron from mammalian serum iron transport proteins (Brown et al., 2010). Interestingly, we previously showed that iron starvation induces expression of virulence factors such as type IV secretion genes (Moumene et al., 2017).

In this study, we show that ERGA_CDS_01230 (UniProt Q5FFA9, named herein Ape for <u>A</u>dhesion <u>p</u>rotein of *<u>E</u>hrlichia ruminantium*,) is involved in the binding of *E. ruminantium* to bovine aortic endothelial cells (BAEC). In order to study whether Ape alone can mediate the invasion of host cells by adhering to endothelial cells membrane, we used latex beads coated with recombinant Ape. These beads covered with the protein adhered and seemed to enter endothelial cells similarly to what is observed with *E. ruminantium in vitro* (Moumene and Meyer, 2015). Subsequent investigation uncovered that rApe is a glycosylated protein that induces a strong humoral immune response in vaccinated goats making it a possible vaccine candidate.

## Methods

### Synchronous culture

*E. ruminantium* (strain Gardel) was propagated in Bovine Aortic endothelial cells (BAEC isolated at CIRAD, Guadeloupe, France) in Baby Hamster Kidney (BHK-21) cell medium supplemented with 1% L-glutamine 200mM (Eurobio), 10% heat-inactivated fetal bovine serum (FBS, Thermofisher), 1% penicillin 10000UI/ streptomycin 10000 μg (Eurobio), 1x amphotericin B (Sigma), 5% NaHCO3 5.5%, under 5% CO2 at 37°C modified from Marcelino et al., 2005. The cell trypsinization (Trypsin-Versen, Eurobio) was carried out with a splitting ratio of ½ when monolayer reached 100% confluence. Synchronous infection of a new BAEC monolayer was obtained using a bacterial suspension previously harvested at 120 hpi by scraping the TC flask (TCF) and passing infected lysed cells through 18G and 26G needles before reinfection at a ratio of 1/20 (Marcelino et al., 2005).

### Quantitative reverse transcription RT-PCR along bacterial developmental cycle

*E. ruminantium*-infected BAE cells were harvested for RNA extraction by trypsinization at 24, 48, 72 and 96 hours post infection (hpi), centrifuged 1700 x g, 5 min at 4°C and by cell lysate scrapping at 120 hpi, centrifuged 20000 x g, 15 min at 4°C. All pellets were dissolved in 1 mL TRIZOL (Thermofisher), RNA was extracted following manufacturer’s recommendations and eluted in 100 μL H2O RNase/DNase free. RNA was treated by Turbo DNAse (Ambion) according to supplier’s instructions and precipitated overnight at −20°C in 2.5 volume (v/v) cold absolute ethanol (Normapur), 1/10 volume of 3 M sodium acetate and 1 μL glycogen 10 mg/mL (Thermofisher). Pellet obtained by centrifugation 15000 x g, 10 min, 4°C was washed with 1 mL 75% ethanol, air dried after centrifugation 9000 x g, 7 min, 4°C and dissolved in 20 μL H2O RNase/ DNase free. Two μg ARN were reverse transcribed using “SuperScript™ VILO™ cDNA Synthesis Kit” (Invitrogen), according to the supplier’s specifications.

### Pre-amplification of *ape* gene

Quantification started by 15 cycles of pre-amplification (same reaction mix and cycling conditions as below). ERGA_CDS_01230 (Ape) was amplified in 25 μL reaction mix containing 250 nM of the forward ERGA_CDS_01230F AATGGAGAATGAGGGGGAAG and reverse ERGA_CDS_01230R ACCCAAACCAAAATCCATCA primers, 12.5 μL of Power SYBR® Green PCR Master Mix (Thermofisher), 12.5 μL H2O RNase/ DNase free and 20 μL pre-amplified DNA. Reactions were performed in a Quantstudio 5 (Thermofisher) as follow: 50°C, 2 min for Uracil-N-Glycosylase activation, 95°C, 10 min for Uracil-N-Glycosylase inactivation and polymerase activation; 40 cycles 95°C, 15 sec denaturation and 59°C, 60 sec hybridization and elongation. The specificity of the PCR product was confirmed by the dissociation curves.

### Quantification of *E. ruminantium* along the bacterial developmental cycle

*E. ruminantium* quantification for normalization as described by (Pruneau et al., 2012), was performed by DNA extraction according to manufacturer’s specifications (QiaAmp DNA minikit, QIAGEN, Courtaboeuf) on 1/10 of the harvested volume, after 20000 x g, 15 min centrifugation and dissolution in 200 μL PBS 1x, followed by a quantitative polymerase chain reaction (qPCR) Sol1 targeting the pCS20 region (Cangi et al., 2017).

### Normalization of *ape* gene expression

Normalization by *E. ruminantium* quantification was calculated: *Rx hpi*= [*cDNA copy number*_(*ERGA_CDS_01230*)_]/[*E. ruminantium pCS20 DNA copy number*], allowing fold change (FC) determination, compared to expression at 96 hpi (corresponding to the stationary phase of the bacterial growth): FC = R _x hpi_/ R_96 hpi_. Results were represented in log2, according to Pruneau et al. (2012) and confirmed by 2 others biological replicates (data not shown).

### Recombinant protein production

pLEXSY_I-blecherry3 plasmid (Jena Bioscience) and 400 ng of amplified CDS_ERGA_01230 were digested 10 min at 37°C by BamHI and SalI (Thermo Scientific, USA) and column purified (Macherey Nagel, Germany). The digested product was ligated in pLEXSY_I-blecherry3 plasmid using T4 DNA Ligase (Thermo Scientific) for 1 h at 22°C. pLEXSY_I-blecherry3 plasmid was modified with the addition of a sequence coding GFP at C-terminus of the insert and 6X His tags at N-terminus of insert. It is to note that, GFP and the coding sequence of CDS_ERGA_01230 were linked with the sequence encoding cleavage site for Xa factor. Ligated plasmid was column purified and 50 ng of the ligation mix was electro-transferred to competent bacteria *E. coli* XL10 (Miller and Nickoloff, 1995). 100 μL of transformed culture were spread on LB medium Petri dish supplemented with carbenicillin (50μg/ml). To confirm the presence of the plasmid and the insert, a PCR was performed on a colony using vector specific primers (Forward, CGCATCACCATCACCATCACG; Reverse, ACCAAAATTGGGACAACACCAGTG). PCR product was then sequenced. Transformed E. coli clone was grown 16 h at 37°C under 200 rpm stirring until optical density (OD) reached 3. The plasmid was isolated using “GenElute™ Plasmid Midiprep” kit (Sigma Aldrich) and digested with *SmaI* (Thermo Scientific)

*Leishmania tarentolae* preculture was grown in 10 mL LEXSY BHI medium with TC flask at 26°C. 3 days after, preculture was diluted 10 fold in 10 mL LEXSY BHI medium and incubated overnight at 26°C (flat position). Next day, *Leishmania tarentolae* were centrifuged 5 min (2000 x g) and half of the medium was removed. Cells were resuspended in remaining medium to get 10^8^ cells/ mL and incubated 10 min on ice. 350 μL cells were electroporated at 450 V, 450 μF, 5 - 6 msec impulsions with 5 μg linearized plasmid (*SmaI* digested). Cells were immediately incubated for 10 min on ice, transfer to 10 mL LEXSY BHI medium (Jena Bioscience) and incubated overnight. 10 μL bleomycin (100 mg/mL, Jena Bioscience) was then added to nonclonal selection for 3 more days. 5 mL of culture supernatant was centrifuged for 3 min at 1000 x g. Pellet was resuspended in 10 mL LEXSY BHI medium containing bleomycin and incubated at 26°C for 5 days. After nonclonal selection, expression of protein was induced in 45 mL BHI medium supplemented with bleomycin and tetracyclin 10 mg/mL (Jena Bioscience). 5 ml of culture from nonclonal selection was added incubated for 72 h at 26°C with shaking at 100 rpm. Supernatant was harvested and concentrated in dialyses bag (3.5 kDa membrane, Serva) in a polyethylene glycol solution 20000 overnight at 4°C. Proteins were concentrated and purified by Sephadex gel filtration.

### Mass spectrometry analysis

Protein size was verified by MALDI-TOF (Microflex, Bruker). The presence of GFP tag and associated fluorescence was checked respectively by immunoblotting and aggregation on Probond beads followed by fluorescence microscopy. Mass spectrometry allowed to check rApe size and Orbitrap allowed to confirm identity of the protein.

### 3D Structure and localization prediction

Protein structure prediction was accomplished using I-TASSER-MTD (Xiaogen Zhou, 2022) and view was generated by MacPyMol (DeLano, 2002). The subcellular localization was predicted by “CELLO 2.5: subCELlular LOcalization predictor” (Yu et al., 2006), from the protein sequence, accession number CAI27575.1. Ape structural homology was determined using “Swiss model, Expasy” (Waterhouse et al., 2018).

### Western Blot for O-GlcNac Glycoprotein detection

rApe was migrated on an acrylamide gel “SDS page” (NuPAGE bis-tris, Novex) for 2 h 30 min at 100V and 400mA in MOPS buffer (Novex), according to the supplier’s instructions (Nu PAGE Technical Guide). Approximately 10 μg protein and BSA for the negative control were denatured at 70°C for 10 min with LDS buffer and reducing agent (Novex) and migrated with a 3 to 198 kDa molecular weight marker (SeeBlue plus 2, Invitrogen). A gel-sized PVDF membrane (Amersham, Hybond-P) was soaked for 30 sec in methanol (Normapur) and incubated in transfer buffer (20x, NuPAGE) for at least 15 min with the same size filter papers (Whatman) and 4 sponges. The transfer assembly was performed according to the NuPAGE technical guide and the transfer was run 1 h 15 min at 30 V, 170 mA. The membrane was then immersed in a solution of Culvert Red (AMRESCO) ~1 min and rinsed with water prior to picture. The protocol for western blot detection was modified from Marcelino et al., 2019.The membrane was blocked for 3 h at room temperature (RT) with stirring in PBS (pH 7.4), 0.05% Tween20 (PBS-T) and 5% milk. Then, membrane was incubated overnight with anti-O-GlcNAc antibody (Santa Cruz), monoclonal IgM, diluted 200 fold in blocking solution. After three washes with PBS-T, membrane was incubated with anti-mouse antibody (IgM-HRP, Molecular probes) diluted 1000 fold in PBS-T for one hour. The membrane was washed 3 times 10 min with PBS-T before the addition of the TMB substrate (Pierce) and gel reader picture once the color was developed.

### Glycosaminoglycans degradation assays

10, 5, 1 and 0.3 μg/ mL heparan sulfate (Jonquieres et al., 2001; Kobayashi et al., 2010) and 9 μg/ mL rApe (positive control) were adsorbed in 50 μL of carbonate/bicarbonate buffer pH 9.6 (Martinez et al., 1993), distributed in Nunc Maxisorp wells, 1 h at 37°C with gentle stirring then overnight at 4°C. The next day, 3 washes were carried out with 200 μL of PBS-T per well. Blocking was done 1 h at 37°C under agitation with 100 μL of blocking buffer PBS tween 20 0.05% milk 3%, followed by 3 washes with 200 μL of PBS-T per well. 50 μL/ well of rApe at 14.3 μg/mL diluted in PBS tween 20 0.05% milk 3% was incubated 1 h at 37°C with stirring. Three washes of 200 μL PBS-T per well were performed, followed by addition of 50 μL anti-GFP antibody diluted to 4,000 in PBS-T with milk 3% and incubation 1 h at 37°C with stirring. Washings were repeated as above. Addition of 200 μL TMB substrate allowed revelation within 30 min at 37°C before to stop the reaction with 100 μL of H2SO4 2N. The reading was done at 450 nm (Multiskan, Thermofisher). 2 cm^2^ wells were inoculated with ~1.1×10^4^ BAEC in 500 μL BHK21 medium. When confluence was reached, several concentrations of chondroitinase (Sigma Aldrich) were tested as follows: 0.2U, 0.4U and 0.9U/ mL chondroitinase was incubated 2 h before infection with the BAEC in 1X PBS and bovine fetal serum (SBF)- free BHK21 medium (Sava et al., 2009; Rajas et al., 2017). The medium was renewed with standard BKH21 prior infection at a ratio 1/20 and 2 h after infection. At lysis stage, all the wells were scraped, centrifuged for 15 min at 20,000 x g. DNA was extracted using the QiaAmp DNA minikit (Qiagen) and quantified using qPCR™ Sol1, targeting the pCS20 region (Cangi et al., 2017). The results were treated using the ΔΔCt method and represented in 2^-ΔΔCt^.

### Flow cytometry for attachment quantification

We used the cytometer to quantify the fluorescence-labelled cells following incubation with rApe. Six-well Nunc plates (9.6 cm^2^/ well) with confluent BAEC were incubated for 2 h at 37°C, 5% CO2 with rApe concentrations ranging from 6.4 to 102.4 μg/ mL following the principle described in (Lundberg et al., 2003). The negative control consisted of confluent BAE cells. After incubation with recombinant protein, each well was rinsed twice with 1 mL of 1X PBS and 1.5 mL of 1X PBS was used to scrape the well. After centrifugation during 10 min at 200 x g, at 4°C cells were resuspended in Isoflow (Beckman) for further reading the percentage of fluorescent labelled cells on the cytometer (FC500, Beckman Coulter).

### Far Western Blot

The BHK21 culture medium of a 175 cm^2^ TCF was removed, the TCF was washed with 5mL of PBS 1x containing anti-protease (Roche). ~10mL of cold PBS 1x was added to gently scrap the cell mat. Centrifugation 10 min at 200 x g and 4°C was performed to remove the supernatant. Lysis of the pellet was performed by addition of 3 mL native lysis buffer (150 mM NaCl, 50 mM Tris-HCl, pH 7.4, NP-40 1%, anti-proteases 1%), followed by 4 “freeze/thaw” cycles by first immersing the lysate 2 min in an ethanol/ ice bath, then 2 min in a 37°C water bath. The lysate was vortexed between each bath. The cells were broken by passing the lysate 4 times through an 18G needle with a syringe. The cell debris were pelleted at 10000 x g for 30 min prior to supernatant transfer into a new tube.

Four different fractions (F1, cytosol components; F2, membrane and organelle components; F3, nucleus components; F4, the cytoskeleton) were extracted using the “ProteoExtract Subcellular Proteome Extraction kit” (Calbiochem/ MERCK) according to the supplier’s instructions. Each fraction was placed in acetone (at least 5x the volume of the extracted fraction) for overnight precipitation at −20°C. The day after, each pellet was re-suspended in 500μL of native lysis buffer after 20 min centrifugation at 15000 x g and 4°C.

18 μL of each cell fraction and 35 μL of lysate were migrated onto acrylamide gel, with addition of LDS, under the same conditions as for the glycosylation tests (but without addition of reducing agent or heating), as well as for the transfer onto PVDF membrane.

The membrane was stored in TTBS 1x (10mM TRIS, 150mM NaCl, 0.05% Tween 20, pH 8,3), then blocked for 1 h at RT under gentle agitation in 2% Bovine Serum Albumine (BSA) in TTBS and finally rinsed 3 times for 5 min with TTBS. The membrane was incubated overnight at 4°C under slow rotation with 0.5 mg rApe in 1% BSA in TTBS. The negative control (without recombinant protein) was incubated in the same conditions. The next day, the membrane was washed 3x for 5 min with TTBS and incubated 1 h at RT under agitation with anti-GFP-HRP (Thermofisher) diluted 2,500 fold in 1% BSA in TTBS. The membrane was washed again 3 x for 5 min with TTBS. The binding of anti-GFP-HRP was revealed by the addition of 4.5 mL peroxide substrate (Pierce Thermofisher) + 0.5mL chromogen DAB (Thermofisher) and incubation 10 min at RT. Picture was taken by colorimetry reading.

### Scanning electronic microscopy for binding assays

Two kinds of fluorescent latex beads (with a sulfate group) were adsorbed with rApe through electrostatic interactions based on (Martinez et al., 2014). A total of 10^10^ beads (0.1 μm in diameter) and 7×10^9^ beads (0.5 μm in diameter) were adsorbed with 100 μg/mL rApe in 1 mL of 25 mM MES, pH 6.1 (Sigma Aldrich) during 4h at RT under slow rotation. Then, three washes were performed using the same buffer before to be resuspended in 1 mL of 1% BSA MES. 24-well plates containing 13 mm diameter lamellae (VWR) were inoculated with BAEC. A deposit of 0.1 or 0.5 μm-diameter beads (1.82×10^8^ per well) was made in BHK21 medium. The negative control consisted of recombinant protein-free beads incubated with BAEC. The plate was centrifuged 5 min at 200 x g at RT before incubation 30-120 min at 37°C under 5% CO2 atmosphere. Three washes with PBS 1X removed the excess of unbound beads before overnight fixation with 2% paraformaldehyde. Lamellae were then removed from each well and dehydrated in series of acetone solutions of increasing concentration, dried to critical point in CO2 and sputter-coated with gold before observation with a FEI Quanta 250 electron microscope at 20 kV.

### ELISA-based binding assays

The antibody response to Ape during vaccination of goats was tested by ELISA. Sera for vaccinated goats obtained from previous studies (Marcelino et al. 2015 a, b) were incubated in wells coated with rApe, followed by incubation with anti-goat IgG antibody coupled to HRP.

The adsorption of 4 μg/ mL rApe diluted in 100 μL of carbonate bicarbonate buffer pH 9.6 was performed in a Nunc plate (Maxisorp). One hour incubation was carried out at 37°C with stirring at 150 rpm then overnight at 4°C. The plate was washed with 300 μL/ well of wash buffer (PBS 1x pH 7.2, Tween 20 0.1%). Blocking was carried out at 37°C with stirring in 300 μL blocking buffer (PBS1x, tween 20 0.1%, casein 2%) for 1 h. Washings were repeated as described in the previous step. 100 μL of each goat serum diluted 100-fold in blocking buffer was incubated 1 h at 37°C with agitation (Perez et al., 1998); 2 blank wells were incubated with 100 μL blocking buffer. Five washes with Wash Buffer preceded the deposit of 100 μL of anti-goat IgG antibody (Rockland) diluted 20,000-fold in blocking buffer and 1 h incubation at 37°C with agitation. Five washes were performed. The revelation was performed by addition of 100 μL of TMB (Neogen) and stopping of the reaction after 5 min development by the addition of 50 μL of 0.5 M H2SO4. Antibody response was detected by ELISA titers and optical density (OD) was read with a spectrophotometer at 450nm. The OD of the wells without serum were valid when < 0.1; OD of negative samples were valid when < 0.2.

## Results

### *ERGA_CDS_01230 (ape*) is highly expressed at infectious elementary body stages of *E. ruminantium* development inside mammalian cells

In order to measure the expression of the *ape* gene, normalization was carried out in relation to the number of bacteria present at each stage of development since no reference gene with a sufficiently stable expression is available for *E. ruminantium*. The developmental cycle of *E. ruminantium* is synchronized when the lysis occurs 5 days after BAEC infection. Quantification of the number of bacteria present in the BAEC every 24 hpi by qPCR Sol1 showed a sigmoidal curve represented in log10 (Figure 1A). The bacterium had a slow growth phase between 24 and 48 hpi then an exponential development with a slowing down of the growth, a stationary phase after 96 hpi and a maximum of copies reached at 120 h (release of elementary bodies). The number of transcripts was determined by qPCR and a ratio calculated at each time as follow: (*E. ruminantium* _cDNA number_) / (*E. ruminantium* _pCS20 DNA copy number_). The fold change (FC) was determined in relation to bacterial expression at 96 hpi (stationnary phase) and represented graphically in Figure 1B. The expression of *ape* gene peaks at the elementary body infectious stages of *E. ruminantium* developmental cycle which correspond to 120 hpi (host cell lysis) and 24 hpi (lag phase). These data indicate that *ape* is expressed when the bacterium is released and ready to infect new cells.

**Figure 1.**
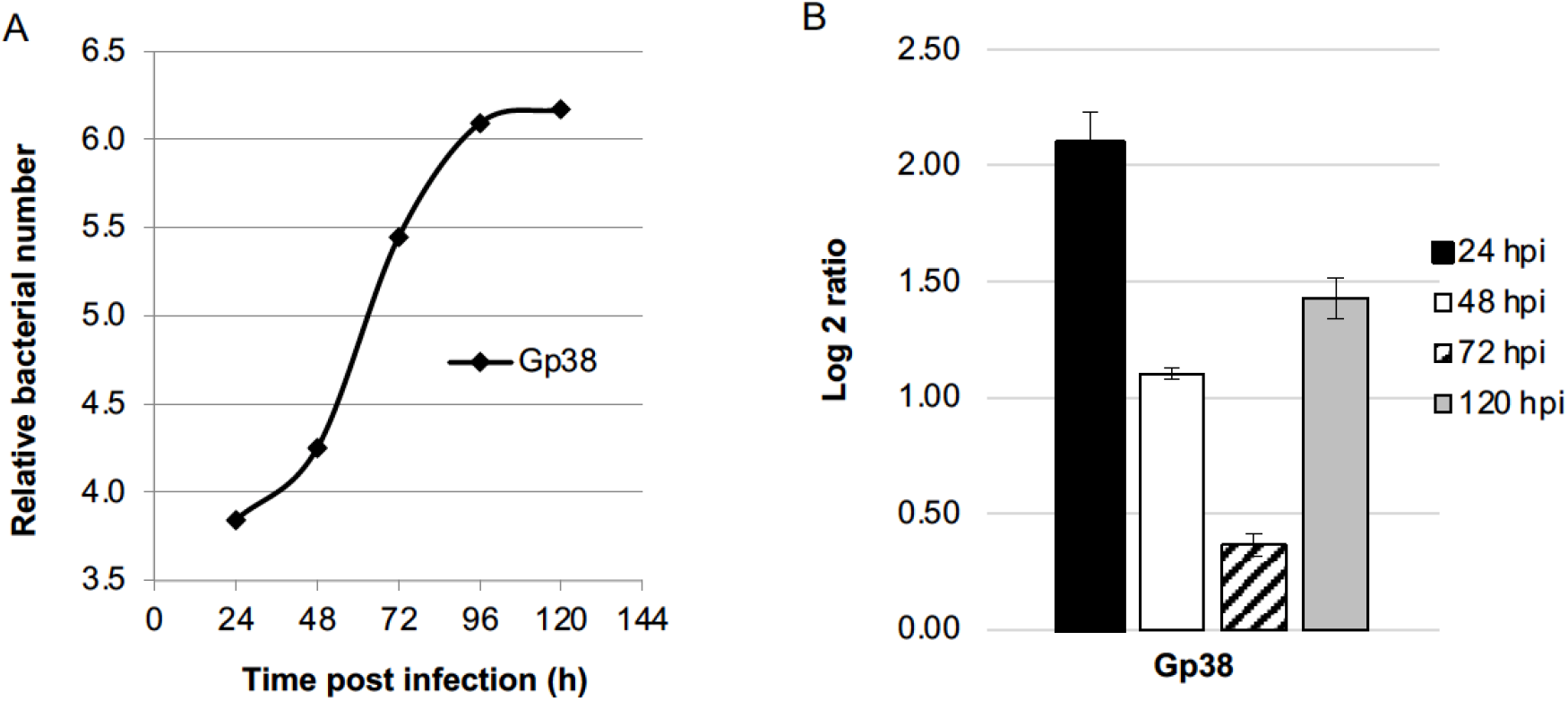
Temporal expression of the (ERGA_CDS_01230) ape gene in *E. ruminantium* determined by qRTPCR. (A) *E. ruminantium* sigmoïdal growth curve was determined by qPCR targeting *pCS20* region and represented as relative bacterial number along the developmental cycle. (B) Transcript levels were determined from 24, 48, 72 and 120 hpi by qRTPCR targeting *ERGA_CDS_01230* gene. Levels were normalized by the quantity of bacteria measured by qPCR Sol1 and the ratio was compared to 96 hpi, allowing fold-change (FC) determination, expressed as log 2. Values at each time point are the means +/- standard deviations for 2 biological replicates. Gp38 is for *E. ruminantium* Gardel strain, passage #38 (virulent strain).

### Ape protein presents a C-clamp structure and is predicted to be an outer-membrane protein

The 3D structure proposed by I-TASSER-MTD software (Figure 2A) showed that Ape harbours a succession of helixes α linked by loops distributed around a β sheet and showed a “c-clamp” three-dimensional structure. By homology with other known I^ry^, II^ry^ or III^ry^ structures, “Swiss model” showed a strong analogy of Ape with a “C-clamp” structure, capable of binding iron. According to the “CELLO 2.5” software, Ape has a dominant cytoplasmic localization (score of 2.418) but is also present at the outer membrane (1.652) (Figure 2B). This canonical sub-cellular localization of a transmembrane protein is in accordance with previous results finding this protein in the outer membrane proteome (Moumene et al., 2015).

**Figure 2.**
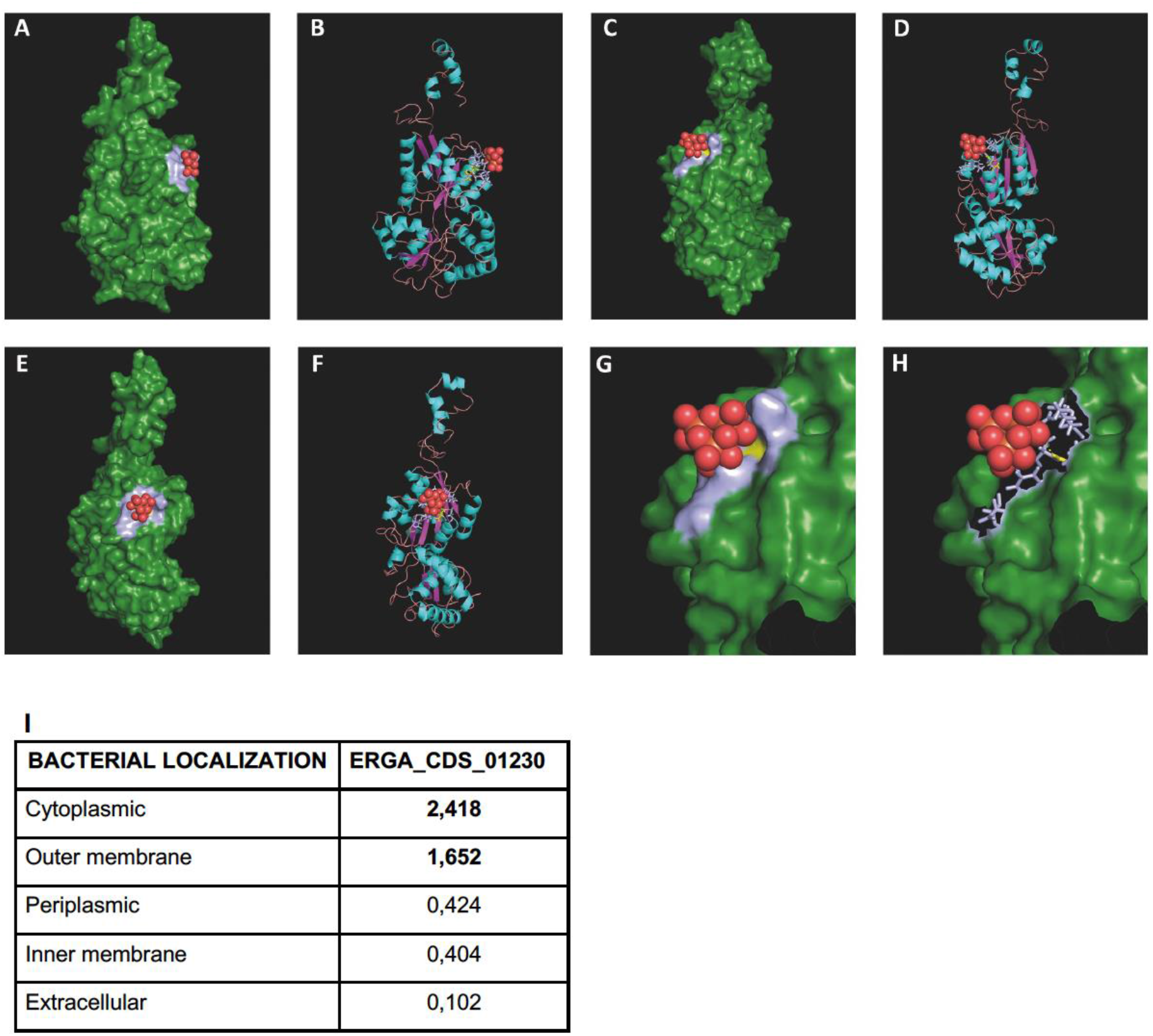
Structural characterization of Ape protein. (A-H) I-TASSER-MTD derived 3D-prediction of the ERGA_CDS_01230 gene product (Ape protein in green) interacting with oxo-iron molecule (red). Front (A-B), back (C-D) and side (E-F) views of Ape showing a succession of α-helixes (turquoise) linked by loops (pink) distributed around β-sheets (purple), organized as a three-dimensional “c-clamp” structure. The enlarged panels (G-H) of the binding site allow the visualization of the residues involved in polar (yellow) and non polar (light blue) interactions. The oxo-iron molecule (CIN1) constituted by 3 atoms of iron (orange) and 12 atoms of oxygen (red) interacts with Ape protein (green) via polar interaction (hydrogen bond with T99 in yellow) and non-polar interactions (hydrophobic interactions in light blue involving L120 and Q122) bonds. (I) Reliability score prediction by CELLO 2.5 software of the subcellular localization of the native *E. ruminantium* Ape protein. Ape presents a typical subcellular localization of an active transporter, with a dominant cytoplasmic localization and is also present at the outer membrane.

### rApe protein shows O-glycosylated post-traductional modifications

Western blot analysis showed that rApe was detected with an anti-O-GlcNAc antibody at the expected size of 66kDa (Figure 3). This size confirmed the data obtained by mass spectrometry and is 30% larger than the one estimated by the amino acid sequence encoded by *ape* gene (41 kDa for 365 amino acids). The identity of rApe protein sequence blasted with the possible major ferric iron binding protein precursor (Q5FFA9). Altogether these results demonstrate the O-glycosylation of rApe.

**Figure 3.**
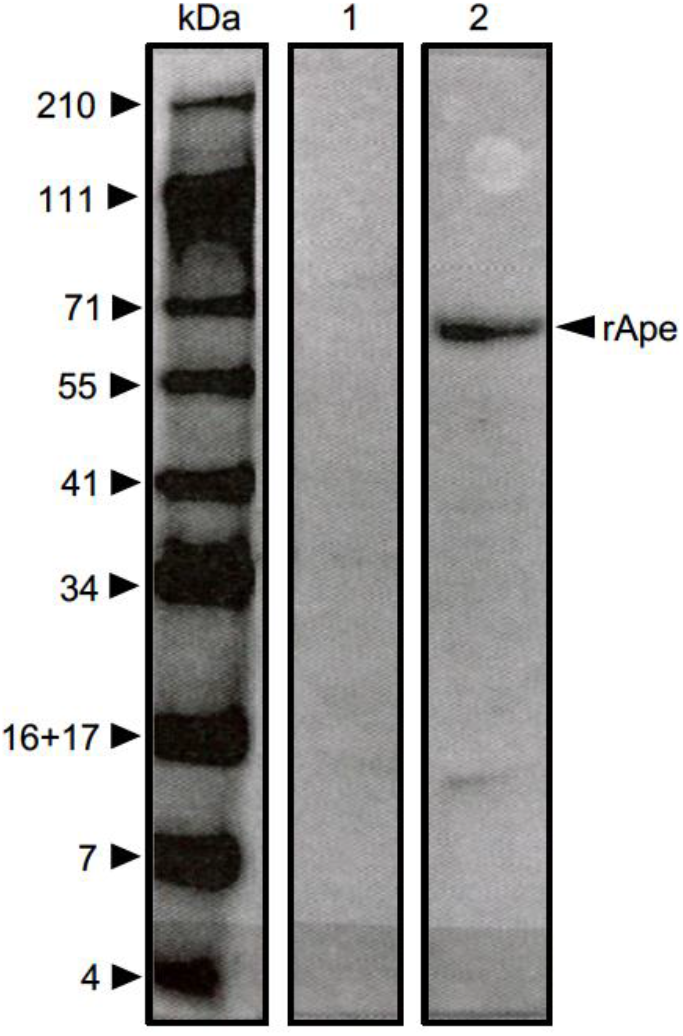
rApe is an O-glycosylated recombinant protein. Composite picture of a Western blot detecting O-glycosylation of recombinant Ape protein (lane 2). Recombinant proteins were separated by SDS-PAGE, then transferred to PVDF and incubated with anti-O-GlcNAc antibody. The Western blot was probed with anti-mouse IgM-HRP antibody and revealed by TMB substrate. Lane 1: negative control: BSA; lane 2: rApe. Numbers and black arrowheads indicate molecular masses in kilodaltons (kDa). The recognized rApe is significantly larger (66 kDa) than the one predicted by the amino acid sequences encoded by ERGA_CDS_01230 gene (41 kDa).

### Enzymatic treatment of BAEC with chondroitinase decreases invasion by *E. ruminantium*

To investigate whether GAGs have a role in *E. ruminantium* adhesion process, BAEC were treated with chondroitinase and then infected with *E. ruminantium*. The number of bacteria was calculated at the end of growth, during the lysis of the BAEC, by qPCR Sol1 for each treatment. After treatment with chondroitinase at 0.2U/mL, the FC corresponding to the bacterial amount differential was higher than for the condition without treatment but this may be explained by inter-well variability (Figure 4). Increasing chondroitinase concentrations to 0.4 and 0.9U/mL resulted in FC of 0.52 and 0.58, respectively, corresponding to a 50% reduction of the number of bacteria compared to the untreated condition, highlighting the role of chondroitin sulfate and dermatan sulfate in adhesion of *E. ruminantium* to the cell. No affinity of rApe for heparan sulfate could by revealed using HRP anti-GFP antibody (data not shown).

**Figure 4.**
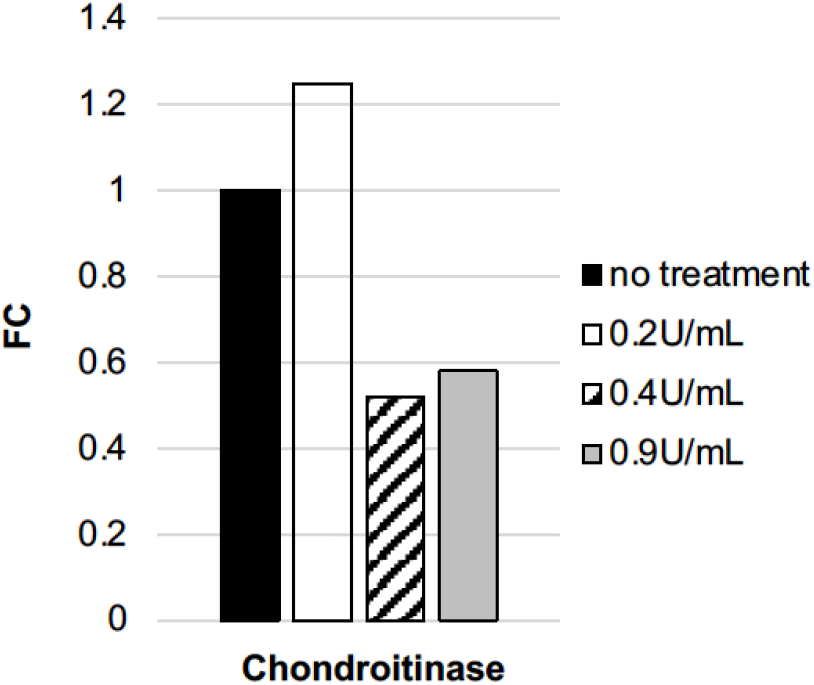
Chondroitinase impairs BAEC infection by *E. ruminantium*. Using 0.4 and 0.9U/mL chondroitinase resulted in a halving of the number of bacteria compared to the untreated condition. Bacterial quantification was performed at lysis stage by qPCR sol1 after chondroitinase treatment at three different concentration. Fold-change (FC) was obtained by calculating the bacterial amount differential for each condition compared to the condition without treatment and represented as 2^-ΔΔCt^.

### rApe binds to BAEC

To go further in the understanding of the role of Ape in *E. ruminantium* invasion of host cells, we analyzed the interaction of rApe with the surface of endothelial cells using flow cytometry. Adhesion of recombinant proteins to BAEC was measured using flow cytometry to detect the fluorescence of rApe harboring GFP tag. The dot plot profile clearly showed that the percentage of labelled BAEC with rApe increases with the amount of rApe protein (Figure 5). As a negative control, the auto-fluorescence of the cells was measured on cells without recombinant protein incubation. The percentage of labelled cells increased with protein concentration up to 36 μg, showing a dose-effect relationship. Above these concentrations, bending of the curve starts with 50% of rApe labelled BAE.

**Figure 5.**
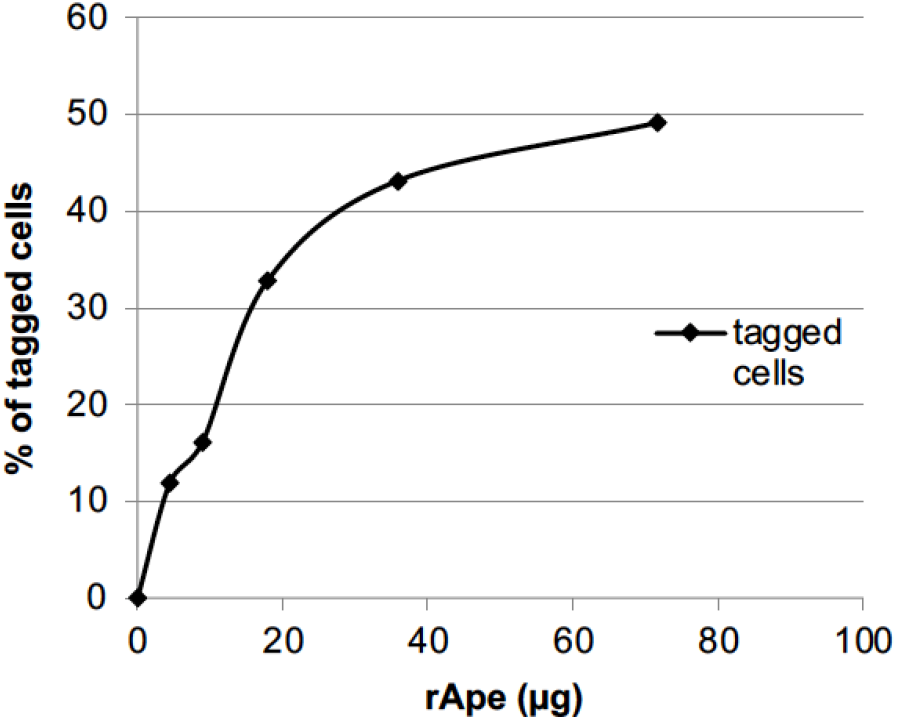
rApe attaches to the host cells. Different concentrations of rApe tagged with GFP were incubated with BAEC. Adherence of GFP-rApe to prefixed BAEC was evaluated by flow cytometry and showed a dose-effect relationship up to 36 μg of rApe. Fluorescent-labelled cells quantified by flow cytometry are represented in % for each concentration point. Auto-fluorescence was evaluated with cells without recombinant protein.

### rApe interacts with BAEC lysate, membrane and organelles cell fractions

The Far Western blot shows the interaction between rApe and the cell fractions as well as the lysate of the BAEC. rApe interacted with proteins of the cell lysate and more specifically with those of the membrane and organelle fraction (Figure 6).

**Figure 6.**
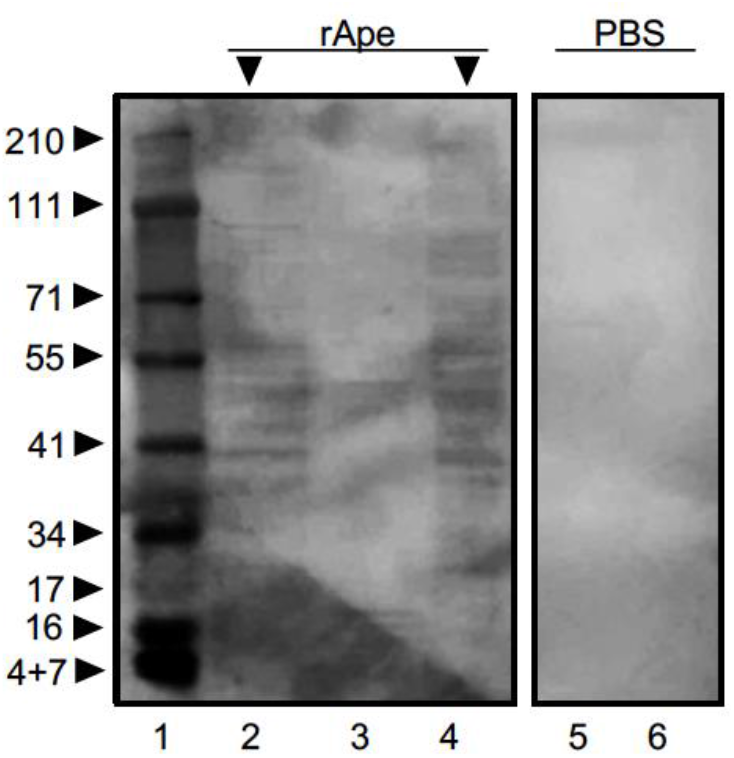
rApe interacts with cell lysate and membrane fractions. Composite picture of a Far-Western blot detecting Ape protein. Lysate and cell fractions were separated by SDS-PAGE, transferred to PVDF and incubated either with rApe (left panel) or PBS as a negative control (right panel). The Western blot was probed with rabbit anti-GFP-HRP antibody, and revealed by peroxide substrate mixed with chromogen DAB. 1: protein ladder, 2 and 5: cell lysates, 3: cytoskeleton fraction, 4 and 6: membrane and organelles fractions.

### rApe-coated beads adhere to BAEC and start internalization mechanism

The visualization of interaction between rApe and the host cell was possible through usage of beads adsorbed with rApe (mimicking *E. ruminantium*) and incubated with BAEC. Adhesion was evaluated by scanning electron microscopy. The negative control showed that non-coated beads did not adhere to the surface of the BAEC, that harbor their classical fried egg shape (Figure 7A). In contrast, Figure 7B showed swollen BAEC, dotted with rApe adsorbed beads after 30 minutes incubation, revealing an interaction between rApe and the BAEC. Figure 7C displayed that rApe-coated beads also adhere (black arrow) and begin to be invaginated (white arrow) by the endothelial cell membrane. Beads diameter did not affect interaction of rApe and the BAEC. The images shown are representative of the observations made in other fields. These data reinforcing the results obtained by immunoblotting and flow cytometry prove that Ape interacts with the host cell membrane and is involved in the adhesion of *E. ruminantium* to the host cell.

**Figure 7.**
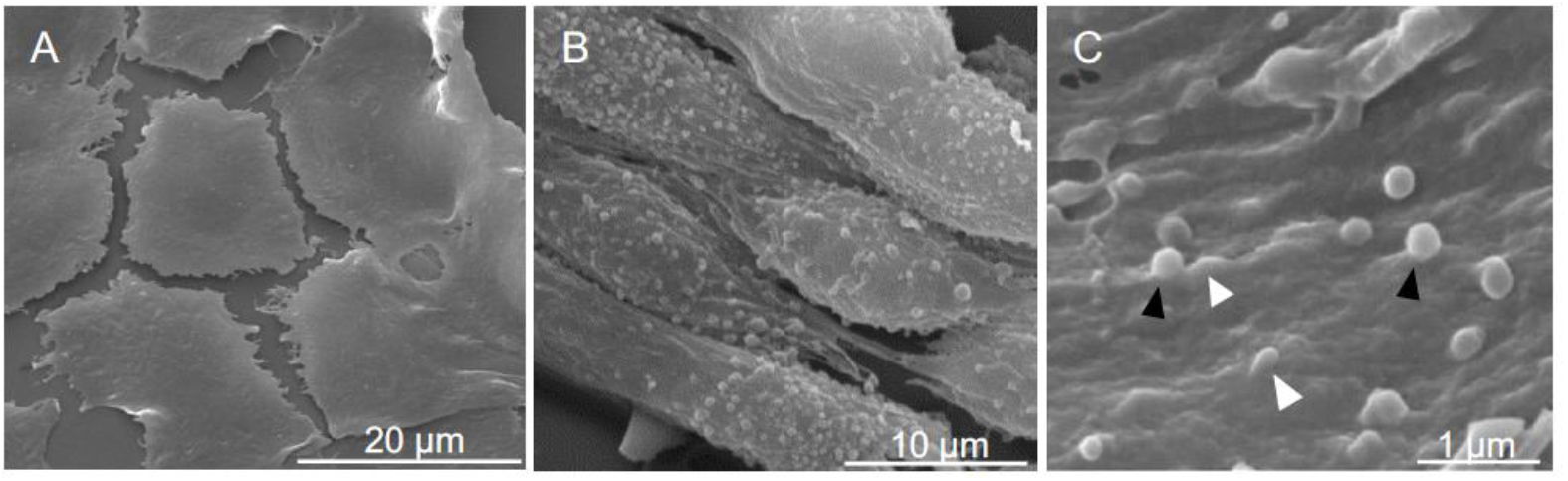
rApe is sufficient for adhesion of latex beads to bovine endothelial cells. Representative images of BAEC incubated 30 minutes with fluorescent latex beads (1.82×10^8^) coated with rApe and processed for scanning electron microscopy. (A) Endothelial cells did not retain non-adsorbed beads (0.1μm diameter) on their cell surface. (B) On the opposite, the whole surface of the same type of cells are covered with adherent beads (0.5μm diameter) absorbed with rApe. (C) Enlarged view of adherent beads. Such beads present two kinds of localizations. Some of them are already internalized (black arrow heads) while others are still located outside the cell remaining in contact with the cytoplasmic membrane (white arrow heads). Scale bars are indicated.

### Ape protein induces an antibody response in vaccinated goats

In order to verify if *E. ruminantium* Ape protein induces a humoral response following vaccination in goats, we tested sera from *in vivo* experiments on animals vaccinated with an inactivated or attenuated bacterial vaccine (Marcelino et al. 2015 a, b). Goats #614 and #915 were both naïve prior to vaccination, characterized by the absence of antibodies against Ape at 3 and 5 weeks post-vaccination, respectively (Figure 8). For #915, the ELISA test showed an increase in humoral response against rApe over time. In fact, the antibody response was developing between 5 and 7 weeks post-vaccination, the latter corresponding to the vaccine boost. For goat #614, inoculation of an attenuated bacterial vaccine also conduced to a humoral response, including response against rApe. These results suggest that rApe could be a relevant target for further studies to see whether it could be a protective immunogen in *E. ruminantium* infection *in vivo*.

**Figure 8.**
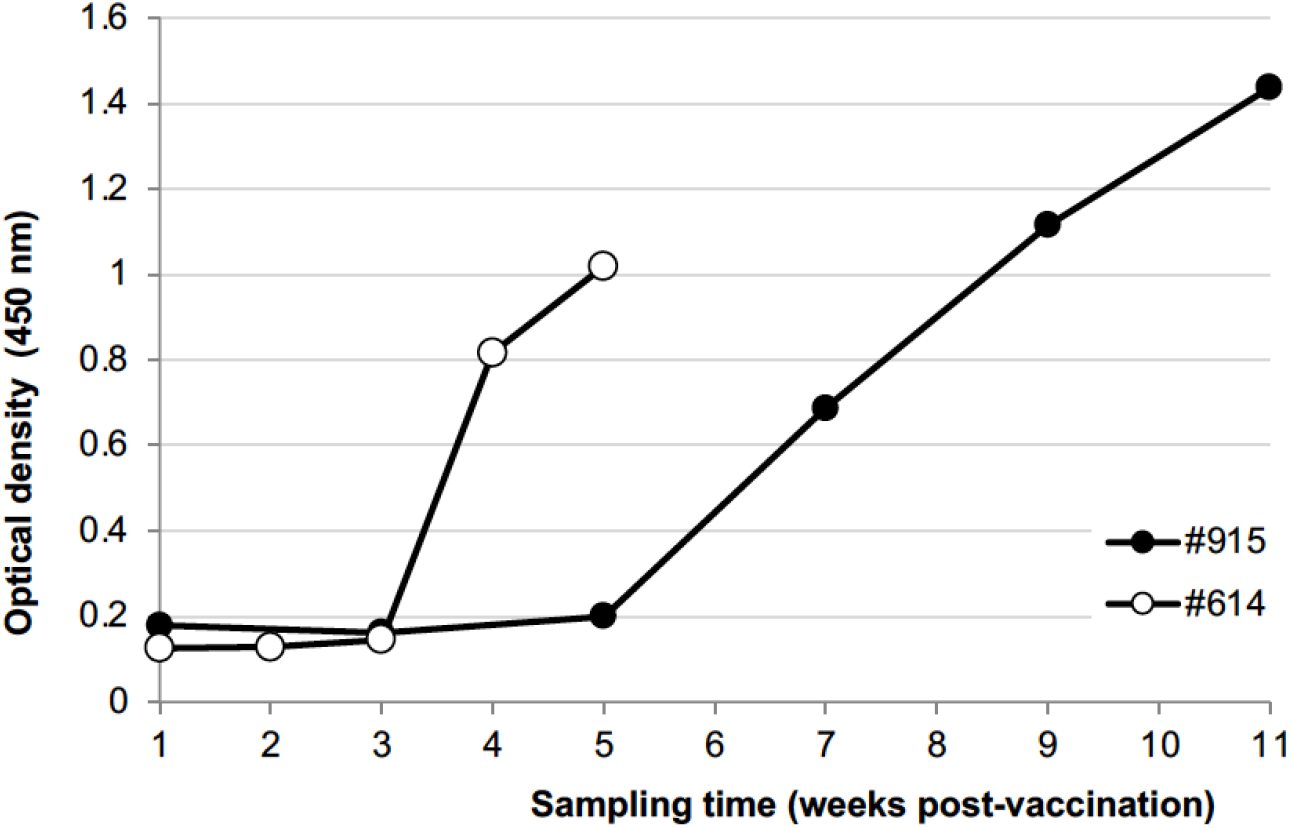
Ape protein induces an antibody response in sera of vaccinated goats. The antibody response to Ape protein during vaccination kinetic was tested by ELISA. Sera of vaccinated goats were incubated in wells coated with rApe, followed by incubation with anti-goat antibody coupled to HRP. Antibody response was detected by ELISA titers (optical density at 450 nm). Shown are representative results from five vaccinated goats. #614: goat vaccinated with an attenuated vaccine. #915: goat vaccinated with an inactivated vaccine. The time (in weeks) post-vaccination is indicated. Goats vaccinated with inactivated vaccine were also challenged for resistance to *E. ruminantium* Gardel strain seven weeks post vaccination.

## Discussion

Controlled and determined *Ehrlichia* entry into host cell is a fundamental first step for an effective infectious developmental cycle, particularly for an obligate intracellular pathogen that strictly relies on its host to grow. In the last decade, comparative genomics and cellular microbiology allowed major discoveries in the molecular pathogenesis of Anaplasmataceae. These integrated approaches led to the effective identification of several bacterial virulence determinants (*e.g*. effectors and regulators) and their diverse mechanisms of action (Martinez et al., 2005; Cheng et al., 2008; Rikihisa, 2017; Marcelino et al., 2015b; Moumene et al., 2017). Notably, Rikihisa *et al*. identified that the EtpE protein governs the binding and entry of *E. chaffeensis* into its host cells (Mohan Kumar et al., 2013). The binding of EtpE to DNaseX elicits a signaling cascade that results in cytoskeleton modification, filopodial induction and finally endocytosis into the host cell (Green et al., 2020). Previous studies demonstrated that functional conservations of molecular pathogenicity determinants can occur between of *E. chaffeensis* and *E. ruminantium* (Moumene et al., 2017) but such bacterial ligand was still unknown in *E. ruminantium* at the beginning of this study. Among other Rickettsiales, a receptor-mediated endocytosis was only reported for *Rickettsia conorii* (Moumene et al., 2017; Martinez et al., 2005).

Our aim being to determine some major pathogenicity determinants of *E. ruminantium*, we chose to analyze ERGA_CDS_01230 (Ape, UniProt Q5FFA9), a putative iron-transporter previously identified in the outer-membrane proteome of *E. ruminantium* (Moumene et al., 2015). Indeed, we postulated that some key bacterial proteins involved in the early interaction with the eukaryotic host cell should be overexpressed at early stages of the developmental cycle and at the bacterial-host interface. Moreover, we previously showed that iron was a triggering environmental cue for several *E. ruminantium* molecular virulence determinants. Indeed, iron depletion induced a master regulatory gene, genes encoding outer-membrane proteins of the Map1 family and genes of the Type IV Secretion System, a major bacterial pathogenicity determinant (Moumene et al., 2017). Therefore, taking into account that iron is a virulence triggering signal of *Ehrlichia*, Ape protein being a putative iron transporter made it an excellent candidate for further characterization. In the present study, we focused on the role of Ape protein in the interaction between *E. ruminantium* and its mammalian host cell, notably during adhesion. As depicted in our working model for *E. ruminantium* binding and invasion of its host endothelial cell (Figure 9), we showed that Ape epitopes are recognized by the immune system of goats vaccinated with live attenuated strain or killed strain or *E. ruminantium*. Indeed, after vaccination, we detected Ape antibodies in sera of vaccinated goats indicating a global humoral response. Following this model, the initial binding of *E. ruminantium* onto the host cell surface seems to involve glycosaminoglycans (GAGs) as chondroitinase treatment of BAEC resulted in a significant decrease of the number of bacteria present at the end of developmental cycle. The *ape* gene is highly expressed at the elementary body developmental stages of *E. ruminantium*, particularly during host cell lysis which precedes *E. ruminantium* release from host cells to initiate a new cycle of infection. Interestingly, recombinant protein rApe is an O-glycosylated protein that interacts with cell membrane and latex beads coated with rApe were able to adhere to the BAEC surface to initiate internalization and follows a similar pattern of entry like that of *E. ruminantium* (Moumene and Meyer, 2015).

**Figure 9.**
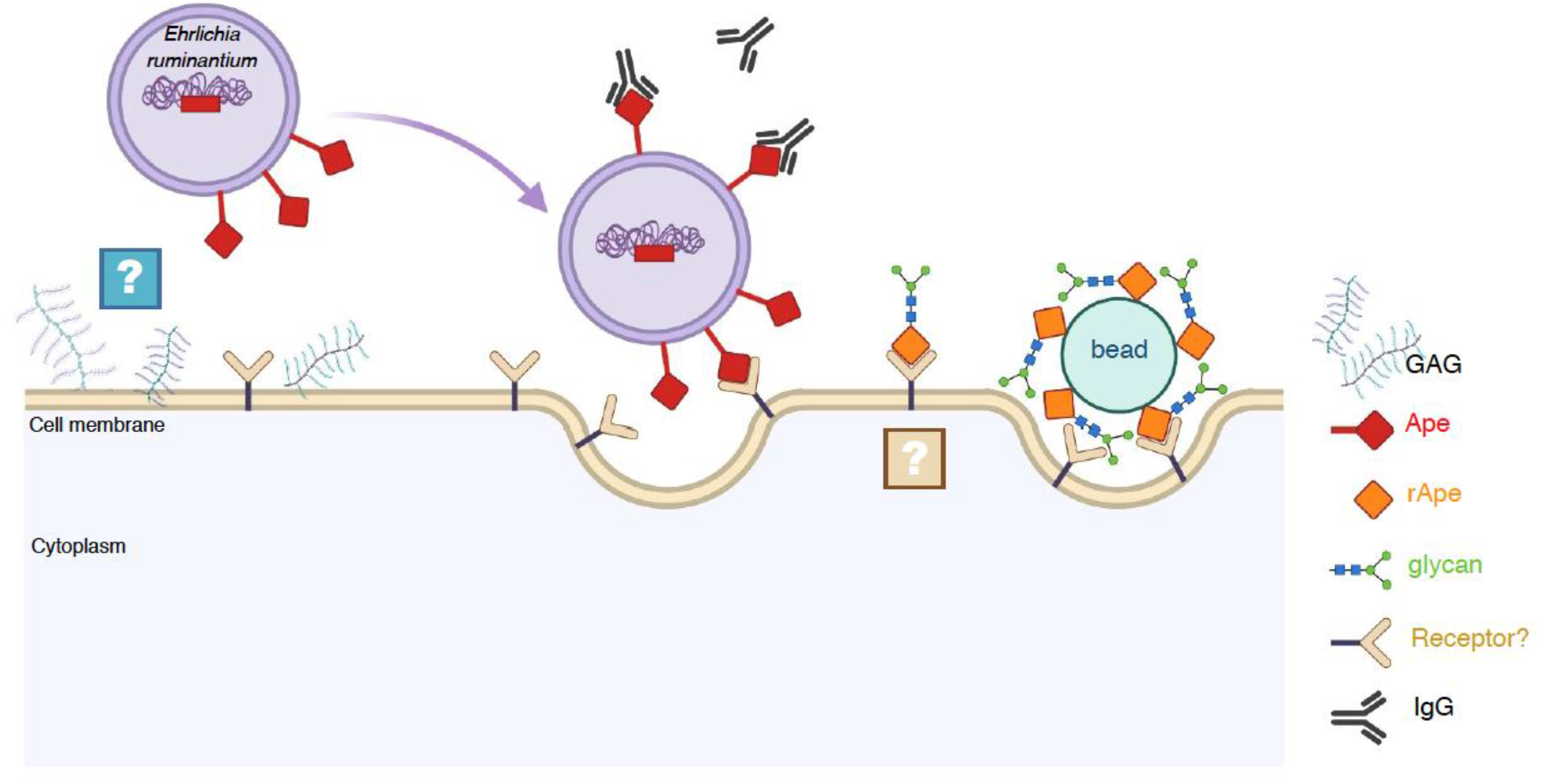
Schematic representation of *Ehrlichia ruminantium* binding to mammalian cells and rApe interaction with the host cell surface. Ape is located at *E. ruminantium* outer membrane and is recognized by antibody from sera of vaccinated animals. *E. ruminantium* can adhere and enter into BAEC but infection is reduced when GAG like chondroitin and dermatan sulfate are degraded. The recombinant version of *E. ruminantium*, rApe, is glycosylated and latex beads coated with rApe bind to BAE cell surface and start to enter in BAEC, in a similar manner that of *E. ruminantium*. Whether Ape binds to a cellular receptor and the following triggered signaling cascade remain to be determined.

Our results showed that Ape protein, an *Ehrlichia* ligand different from previously identified *E. chaffeensis* EtpE (Mohan Kumar et al., 2013), is important for *E. ruminantium* adhesion to the mammalian host cells. The mechanisms used by *E. ruminantium* to invade its host are still not elucidated compared to other pathogens of the order *Rickettsiales*. Indeed *Rickettsia conorii* uses OmpA (Hillman et al., 2013) to adhere to the host cell whereas two different receptors were described for *Anaplasma marginale* and *A. phagocytophilum* mobilizing Msp1a and OmpA (de la Fuente et al., 2003; Hebert et al., 2017), Asp14 and OmpA (Kahlon et al., 2013; Ojogun et al., 2012), respectively to attach cell membrane. Adhesins and invasins were largely described in other bacteria as surface located structures for specific interaction with host cell receptors (Niemann et al., 2004). *Yersinia* outer membrane invasin interacts with ß1 integrin receptors, inducing several reactions including actin rearrangements at the site of bacterial entry, promoting invasion. *Salmonella* translocates several effectors into target cells, some of them allowing the initial uptake of the bacterium, whereas *Listeria* uses InlA and InlB-dependent molecular pathways, (Pizarro-Cerda and Cossart, 2006). Ligands often show elongated molecules containing domains commonly found in eukaryotic proteins (Niemann et al., 2004).

We showed that rApe is O-glycosylated. Post-translational modifications are one of the most important mechanisms for activating, changing, or suppressing protein functions, being widely used by pathogens to interact with their hosts. *E. ruminantium* glycoproteomics showed a high percentage of glycoproteins, many of them being O-glycosylated (Marcelino et al., 2019). *E. ruminantium* “mucin”, which is also glycosylated, was presented as an adhesin for tick cells, reinforcing a role of glycans in *Ehrlichia* adhesin molecules (de la Fuente et al., 2004). The strength of ligand-receptor bacterial interactions is optimized depending on their environment but weak enough to allow a bacterium to detach regularly and migrate to other locations (Formosa-Dague et al., 2018). Glycan-glycan interactions in bacterial-mammalian cells systems were characterized as low-affinity weak interactions preceding high-affinity protein-glycan or protein-protein interactions but recent studies have documented the importance of such interactions in bacterial adhesion (Formosa-Dague et al., 2018). Indeed, we determined that the presence GAG on the surface of BAE plays a key role in the attachment of *Ehrlichia* to bovine endothelial cells *in vitro*, reinforcing the hypothesis that several receptors are probably required in *E. ruminantium* adhesion and subsequent infection of host cells. Chondroitinase treatment significantly affected *Ehrlichia* entry compared to the untreated condition. The enzymatic digestion of chondroitin and dermatan from BAEC reduced the rate of infection of the BAEC, as *E. ruminantium* can no longer adhere to the surface of the cells. Indeed, in other models like *Chlamydia*, GAGs were shown to be used for initial attachment to host cells (Tiwari et al., 2012); Lyme disease *Borreliae* requires glycosaminoglycan binding activity to colonize and disseminate to tissues (Lin et al., 2017). Even though heparan sulfate appears to be the most important GAG species involved in bacterial binding, both heparan sulfate and chondroitin sulfate were able to influence the attachment of mucoid *P. aeruginosa, H. influenza* and *B. cepacia* in specific ways that were dependent on the cell line involved (Martin et al., 2019). *Borrelia burgdorferi* has multiple surface proteins with different binding specificities to GAGs depending on the tissue affected (Leong et al., 1998). Different GAGs act as receptors for *B. burgdorferi* depending on the host cells; both heparin sulfate and heparan sulfate are essential in adherence to primary endothelium and adult kidney Vero cells, but only dermatan sulfate is involved in attachment to human embryonic kidney cells (Garcia et al., 2016).

Although we did not establish the interaction of rApe adhesion with a cellular receptor, we can still hypothesize that Ape protein actively triggers *Ehrlichia* internalization by the mean of a ligand/receptor interaction. Kumar et al. already suggested the existence of additional mammalian receptors for *Ehrlichia* infection (Mohan Kumar et al., 2013). The present study identifies the bacterial ligand of a putative second *Ehrlichia* invasin-receptor pair and highlights the importance of this molecular control of invasion for the *Anaplasmataceae* intracellular bacteria. The probable existence of several ligand-receptor systems could indeed serve the bacteria to infect a broader host range of animal reservoirs and vector ticks. Moreover, GAG degradation following chondroitinase treatment severely impaired *Ehrlichia* infection. This suggests that Ape could interact with a glycosylphosphatidylinositol (GPI)-anchored protein as previously shown for PSGL-1 that is required for the binding and infection of human HL-60 cells by *A. phagocytophilum* (Herron et al., 2000). This remains to be further studied as well as its role in iron uptake of *Ehrlichia ruminantium* (Reneer et al., 2008).

In summary, with the identification of Ape protein (ERGA_CDS_01230), we found the first *Ehrlichia ruminantium* protein that is involved in host cell invasion. Whether the *E. chaffeensis* EtpE homolog (ERGA_CDS_08340) is functional in *E. ruminantium* remain to be explored, but these outer-membrane proteins can now be considered as immune-dominant pathogen-associated molecular patterns (Budachetri et al., 2020). Our next step is now to investigate the use of rApe as a new vaccine candidate against Heartwater. In light of the lack of prophylactic measures against *Ehrlichia* spp. and the rising appearance of antibiotic resistances, deep understanding of invasion mechanisms is of prime importance and will help to propose efficient alternative therapeutics blocking the early interaction between these obligate intracellular bacteria and their host cells.

## Supporting information

RawData

Readme File for RawData

## Acknowledgements

The authors gratefully acknowledge Géraldine Bossard and Valérie Rodrigues for technical assistance in the development of ELISA assays. Preprint version 3 of this article has been peer-reviewed and recommended by Peer Community In Infections (https://doi.org/10.24072/pci.infections.100003).

## Data, scripts, code, and supplementary information availability

Data are available online at https://doi.org/10.1101/2021.06.15.447525

## References

Allsopp, B.A. (2010). Natural history of *Ehrlichia ruminantium*. Vet Parasitol, 167, 123–135. https://doi.org/10.1016/j.vetpar.2009.09.014

Aquino, R.S., Lee, E.S., and Park, P.W. (2010). Diverse functions of glycosaminoglycans in infectious diseases. Prog Mol Biol Transl Sci, 93, 373–394. https://doi.org/10.1016/S1877-1173(10)93016-0

Aquino, R.S., Teng, Y.H., and Park, P.W. (2018). Glycobiology of syndecan-1 in bacterial infections. Biochem Soc Trans. https://doi.org/10.1042/BST20170395

Bartlett, A.H., and Park, P.W. (2010). Proteoglycans in host-pathogen interactions: molecular mechanisms and therapeutic implications. Expert Rev Mol Med, 12, e5. https://doi.org/10.1017/S1462399409001367

Brown, R.N., Romine, M.F., Schepmoes, A.A., Smith, R.D., and Lipton, M.S. (2010). Mapping the subcellular proteome of *Shewanella oneidensis* MR-1 using sarkosyl-based fractionation and LC-MS/MS protein identification. J Proteome Res, 9, 4454–4463. https://doi.org/10.1021/pr100215h

Budachetri, K., Teymournejad, O., Lin, M., Yan, Q., Mestres-Villanueva, M., Brock, G.N., and Rikihisa, Y. (2020). An Entry-Triggering Protein of *Ehrlichia* Is a New Vaccine Candidate against Tick-Borne Human Monocytic Ehrlichiosis. mBio, 11. https://doi.org/10.1128/mBio.00895-20

Cangi, N., Gordon, J.L., Bournez, L., Pinarello, V., Aprelon, R., Huber, K., Lefrancois, T., Neves, L., Meyer, D.F., and Vachiery, N. (2016). Recombination Is a Major Driving Force of Genetic Diversity in the *Anaplasmataceae Ehrlichia ruminantium*. Front Cell Infect Microbiol, 6, 111. https://doi.org/10.3389/fcimb.2016.00111

Cangi, N., Pinarello, V., Bournez, L., Lefrancois, T., Albina, E., Neves, L., and Vachiery, N. (2017). Efficient high-throughput molecular method to detect *Ehrlichia ruminantium* in ticks. Parasit Vectors, 10, 566. https://doi.org/10.1186/s13071-017-2490-0

Cheng, Z., Wang, X., and Rikihisa, Y. (2008). Regulation of type IV secretion apparatus genes during *Ehrlichia chaffeensis* intracellular development by a previously unidentified protein. J Bacteriol, 190, 2096–2105. https://doi.org/10.1128/JB.01813-07

De La Fuente, J., Garcia-Garcia, J.C., Barbet, A.F., Blouin, E.F., and Kocan, K.M. (2004). Adhesion of outer membrane proteins containing tandem repeats of *Anaplasma* and *Ehrlichia* species (Rickettsiales: *Anaplasmatacea*e) to tick cells. Vet Microbiol, 98, 313–322. https://doi.org/10.1016/j.vetmic.2003.11.001

De La Fuente, J., Garcia-Garcia, J.C., Blouin, E.F., and Kocan, K.M. (2003). Characterization of the functional domain of major surface protein 1a involved in adhesion of the rickettsia *Anaplasma marginale* to host cells. Vet Microbiol, 91, 265–283. https://doi.org/10.1016/s0378-1135(02)00309-7

Deem, S.L. (1998). A review of heartwater and the threat of introduction of *Cowdria ruminantium* and *Amblyomma* spp. ticks to the American mainland. J Zoo Wildl Med, 29, 109–113.

Warren L. DeLano “The PyMOL Molecular Graphics System.” DeLano Scientific, San Carlos, CA, USA. http://www.pymol.org

Dumler, J.S., Barbet, A.F., Bekker, C.P., Dasch, G.A., Palmer, G.H., Ray, S.C., Rikihisa, Y., and Rurangirwa, F.R. (2001). Reorganization of genera in the families *Rickettsiaceae* and *Anaplasmataceae* in the order Rickettsiales: unification of some species of *Ehrlichia* with *Anaplasma, Cowdria* with *Ehrlichia* and *Ehrlichia* with *Neorickettsia,* descriptions of six new species combinations and designation of *Ehrlichia equi* and ‘HGE agent’ as subjective synonyms of *Ehrlichia phagocytophila*. Int J Syst Evol Microbiol, 51, 2145–2165. https://doi.org/10.1099/00207713-51-6-2145

Formosa-Dague, C., Castelain, M., Martin-Yken, H., Dunker, K., Dague, E., and Sletmoen, M. (2018). The Role of Glycans in Bacterial Adhesion to Mucosal Surfaces: How Can Single-Molecule Techniques Advance Our Understanding? Microorganisms, 6. https://doi.org/10.3390/microorganisms6020039

Frutos, R., Viari, A., Ferraz, C., Bensaid, A., Morgat, A., Boyer, F., Coissac, E., Vachiery, N., Demaille, J., and Martinez, D. (2006). Comparative genomics of three strains of *Ehrlichia ruminantium:* a review. Ann N Y Acad Sci, 1081, 417–433. https://doi.org/10.1196/annals.1373.061

Frutos, R., Viari, A., Vachiery, N., Boyer, F., and Martinez, D. (2007). *Ehrlichia ruminantium:* genomic and evolutionary features. Trends Parasitol, 23, 414–419. https://doi.org/10.1016/j.pt.2007.07.007

Gagoski, D., Mureev, S., Giles, N., Johnston, W., Dahmer-Heath, M., Skalamera, D., Gonda, T.J., and Alexandrov, K. (2015). Gateway-compatible vectors for high-throughput protein expression in pro-and eukaryotic cell-free systems. J Biotechnol, 195, 1–7. https://doi.org/10.1016/j.jbiotec.2014.12.006

Garcia, B., Merayo-Lloves, J., Martin, C., Alcalde, I., Quiros, L.M., and Vazquez, F. (2016). Surface Proteoglycans as Mediators in Bacterial Pathogens Infections. Front Microbiol, 7, 220. https://doi.org/10.3389/fmicb.2016.00220

Green, R.S., Izac, J.R., Naimi, W.A., O’bier, N., Breitschwerdt, E.B., Marconi, R.T., and Carlyon, J.A. (2020). *Ehrlichia chaffeensis* EplA Interaction With Host Cell Protein Disulfide Isomerase Promotes Infection. Front Cell Infect Microbiol, 10, 500. https://doi.org/10.3389/fcimb.2020.00500

Hebert, K.S., Seidman, D., Oki, A.T., Izac, J., Emani, S., Oliver, L.D., Jr., Miller, D.P., Tegels, B.K., Kannagi, R., Marconi, R.T., and Carlyon, J.A. (2017). *Anaplasma marginale* Outer Membrane Protein A Is an Adhesin That Recognizes Sialylated and Fucosylated Glycans and Functionally Depends on an Essential Binding Domain. Infect Immun, 85. https://doi.org/10.1128/IAI.00968-16

Herron, M.J., Nelson, C.M., Larson, J., Snapp, K.R., Kansas, G.S., and Goodman, J.L. (2000). Intracellular parasitism by the human granulocytic ehrlichiosis bacterium through the P-selectin ligand, PSGL-1. Science, 288, 1653–1656. https://doi.org/10.1126/science.288.5471.1653

Hillman, R.D., Baktash, Y.M., and Martinez, J.J. (2013). OmpA-mediated rickettsial adherence to and invasion of human endothelial cells is dependent upon interaction with alpha2beta1 integrin. Cell Microbiol, 15, 727–741. https://doi.org/10.1111/cmi.12068

Jonquieres, R., Pizarro-Cerda, J., and Cossart, P. (2001). Synergy between the N-and C-terminal domains of InlB for efficient invasion of non-phagocytic cells by Listeria monocytogenes. Mol Microbiol, 42, 955–965. https://doi.org/10.1046/j.1365-2958.2001.02704.x

Kahlon, A., Ojogun, N., Ragland, S.A., Seidman, D., Troese, M.J., Ottens, A.K., Mastronunzio, J.E., Truchan, H.K., Walker, N.J., Borjesson, D.L., Fikrig, E., and Carlyon, J.A. (2013). *Anaplasma phagocytophilum* Asp14 is an invasin that interacts with mammalian host cells via its C terminus to facilitate infection. Infect Immun, 81, 65–79. https://doi.org/10.1128/IAI.00932-12

Kobayashi, K., Kato, K., Sugi, T., Takemae, H., Pandey, K., Gong, H., Tohya, Y., and Akashi, H. (2010). *Plasmodium falciparum* BAEBL binds to heparan sulfate proteoglycans on the human erythrocyte surface. J Biol Chem, 285, 1716–1725. https://doi.org/10.1074/jbc.M109.021576

Leong, J.M., Robbins, D., Rosenfeld, L., Lahiri, B., and Parveen, N. (1998). Structural requirements for glycosaminoglycan recognition by the Lyme disease spirochete, *Borrelia burgdorferi*. Infect Immun, 66, 6045–6048. https://doi.org/10.1128/IAI.66.12.6045-6048.1998

Lin, M., Liu, H., Xiong, Q., Niu, H., Cheng, Z., Yamamoto, A., and Rikihisa, Y. (2016). *Ehrlichia* secretes Etf-1 to induce autophagy and capture nutrients for its growth through RAB5 and class III phosphatidylinositol 3-kinase. Autophagy, 12, 2145–2166. https://doi.org/10.1080/15548627.2016.1217369

Lin, Y.P., Li, L., Zhang, F., and Linhardt, R.J. (2017). *Borrelia burgdorferi* glycosaminoglycan-binding proteins: a potential target for new therapeutics against Lyme disease. Microbiology, 163, 1759–1766. https://doi.org/10.1099/mic.0.000571

Lundberg, M., Wikstrom, S., and Johansson, M. (2003). Cell surface adherence and endocytosis of protein transduction domains. Mol Ther, 8, 143–150. https://doi.org/10.1016/s1525-0016(03)00135-7

Marcelino, I., Colome-Calls, N., Holzmuller, P., Lisacek, F., Reynaud, Y., Canals, F., and Vachiery, N. (2019). Sweet and Sour *Ehrlichia:* Glycoproteomics and Phosphoproteomics Reveal New Players in *Ehrlichia ruminantium* Physiology and Pathogenesis. Front Microbiol, 10, 450. https://doi.org/10.3389/fmicb.2019.00450

Marcelino, I., Verissimo, C., Sousa, M.F., Carrondo, M.J., and Alves, P.M. (2005). Characterization of *Ehrlichia ruminantium* replication and release kinetics in endothelial cell cultures. Vet Microbiol, 110, 87–96. https://doi.org/10.1016/j.vetmic.2005.07.012

Martin, C., Lozano-Iturbe, V., Giron, R.M., Vazquez-Espinosa, E., Rodriguez, D., Merayo-Lloves, J., Vazquez, F., Quiros, L.M., and Garcia, B. (2019). Glycosaminoglycans are differentially involved in bacterial binding to healthy and cystic fibrosis lung cells. J Cyst Fibros, 18, e19–e25. https://doi.org/10.1016/j.jcf.2018.10.017

Martinez, D., Coisne, S., Sheikboudou, C., and Jongejan, F. (1993). Detection of antibodies to *Cowdria ruminantium* in the serum of domestic ruminants by indirect ELISA. Rev Elev Med Vet Pays Trop, 46, 115–120.

Martinez, E., Cantet, F., Fava, L., Norville, I., and Bonazzi, M. (2014). Identification of OmpA, a *Coxiella burnetii* protein involved in host cell invasion, by multi-phenotypic high-content screening. PLoS Pathog, 10, e1004013. https://doi.org/10.1371/journal.ppat.1004013

Martinez, J.J., Seveau, S., Veiga, E., Matsuyama, S., and Cossart, P. (2005). Ku70, a component of DNA-dependent protein kinase, is a mammalian receptor for *Rickettsia conorii*. Cell, 123, 1013–1023. https://doi.org/10.1016/j.cell.2005.08.046

Miller, E.M., and Nickoloff, J.A. (1995). *Escherichia coli* electrotransformation. Methods Mol Biol, 47, 105–113. https://doi.org/10.1385/0-89603-310-4:105

Mohan Kumar, D., Yamaguchi, M., Miura, K., Lin, M., Los, M., Coy, J.F., and Rikihisa, Y. (2013). *Ehrlichia chaffeensis* uses its surface protein EtpE to bind GPI-anchored protein DNase X and trigger entry into mammalian cells. PLoS Pathog, 9, e1003666. https://doi.org/10.1371/journal.ppat.1003666

Moumene, A., Gonzalez-Rizzo, S., Lefrancois, T., Vachiery, N., and Meyer, D.F. (2017). Iron Starvation Conditions Upregulate *Ehrlichia ruminantium* Type IV Secretion System, *tr1* Transcription Factor and *map1* Genes Family through the Master Regulatory Protein ErxR. Front Cell Infect Microbiol, 7, 535. https://doi.org/10.3389/fcimb.2017.00535

Moumene, A., Marcelino, I., Ventosa, M., Gros, O., Lefrancois, T., Vachiery, N., Meyer, D.F., and Coelho, A.V. (2015). Proteomic profiling of the outer membrane fraction of the obligate intracellular bacterial pathogen *Ehrlichia ruminantium*. PLoS One, 10, e0116758. https://doi.org/10.1371/journal.pone.0116758

Moumene, A., and Meyer, D.F. (2016). *Ehrlichia’s* molecular tricks to manipulate their host cells. Microbes Infect, 18-3, 172–179. https://doi.org/10.1016/j.micinf.2015.11.001

Niemann, H.H., Schubert, W.D., and Heinz, D.W. (2004). Adhesins and invasins of pathogenic bacteria: a structural view. Microbes Infect, 6, 101–112. https://doi.org/10.1016/j.micinf.2003.11.001

Noroy, C., Lefrancois, T., and Meyer, D.F. (2019). Searching algorithm for Type IV effector proteins (S4TE) 2.0: Improved tools for Type IV effector prediction, analysis and comparison in proteobacteria. PLoS Comput Biol, 15, e1006847. https://doi.org/10.1371/journal.pcbi.1006847

Ojogun, N., Kahlon, A., Ragland, S.A., Troese, M.J., Mastronunzio, J.E., Walker, N.J., Viebrock, L., Thomas, R.J., Borjesson, D.L., Fikrig, E., and Carlyon, J.A. (2012). *Anaplasma phagocytophilum* outer membrane protein A interacts with sialylated glycoproteins to promote infection of mammalian host cells. Infect Immun, 80, 3748–3760. https://doi.org/10.1128/IAI.00654-12

Perez, J.M., Martinez, D., Sheikboudou, C., Jongejan, F., and Bensaid, A. (1998). Characterization of variable immunodominant antigens of *Cowdria ruminantium* by ELISA and immunoblots. Parasite Immunol, 20, 613–622. https://doi.org/10.1046/j.1365-3024.1998.00193.x

Pizarro-Cerda, J., and Cossart, P. (2006). Bacterial adhesion and entry into host cells. Cell, 124, 715–727. https://doi.org/10.1016/j.cell.2006.02.012

Pruneau, L., Emboule, L., Gely, P., Marcelino, I., Mari, B., Pinarello, V., Sheikboudou, C., Martinez, D., Daigle, F., Lefrancois, T., Meyer, D.F., and Vachiery, N. (2012). Global gene expression profiling of *Ehrlichia ruminantium* at different stages of development. FEMS Immunol Med Microbiol, 64, 66–73. https://doi.org/10.1111/j.1574-695X.2011.00901.x

Rajas, O., Quiros, L.M., Ortega, M., Vazquez-Espinosa, E., Merayo-Lloves, J., Vazquez, F., and Garcia, B. (2017). Glycosaminoglycans are involved in bacterial adherence to lung cells. BMC Infect Dis, 17, 319. https://doi.org/10.1186/s12879-017-2418-5

Reneer, D.V., Troese, M.J., Huang, B., Kearns, S.A., and Carlyon, J.A. (2008). *Anaplasma phagocytophilum* PSGL-1-independent infection does not require Syk and leads to less efficient AnkA delivery. Cell Microbiol, 10, 1827–1838. https://doi.org/10.1111/j.1462-5822.2008.01168.x

Rikihisa, Y. (2017). Role and Function of the Type IV Secretion System in *Anaplasma* and *Ehrlichia* Species. Curr Top Microbiol Immunol, 413, 297–321. https://doi.org/10.1007/978-3-319-75241-9_12

Sava, I.G., Zhang, F., Toma, I., Theilacker, C., Li, B., Baumert, T.F., Holst, O., Linhardt, R.J., and Huebner, J. (2009). Novel interactions of glycosaminoglycans and bacterial glycolipids mediate binding of enterococci to human cells. J Biol Chem, 284, 18194–18201. https://doi.org/10.1074/jbc.M901460200

Teymournejad, O., Lin, M., and Rikihisa, Y. (2017). *Ehrlichia chaffeensis* and Its Invasin EtpE Block Reactive Oxygen Species Generation by Macrophages in a DNase X-Dependent Manner. mBio, 8. https://doi.org/10.1128/mBio.01551-17

Tiwari, V., Maus, E., Sigar, I.M., Ramsey, K.H., and Shukla, D. (2012). Role of heparan sulfate in sexually transmitted infections. Glycobiology, 22, 1402–1412. https://doi.org/10.1093/glycob/cws106

Truttmann, M.C., Rhomberg, T.A., and Dehio, C. (2011). Combined action of the type IV secretion effector proteins BepC and BepF promotes invasome formation of *Bartonella henselae* on endothelial and epithelial cells. Cell Microbiol, 13, 284–299. https://doi.org/10.1111/j.1462-5822.2010.01535.x

Waterhouse, A., Bertoni, M., Bienert, S., Studer, G., Tauriello, G., Gumienny, R., Heer, F.T., De Beer, T.a.P., Rempfer, C., Bordoli, L., Lepore, R., and Schwede, T. (2018). SWISS-MODEL: homology modelling of protein structures and complexes. Nucleic Acids Res, 46, W296–W303. https://doi.org/10.1093/nar/gky427

Xiaogen Zhou, W.Z., Yang Li, Robin Pearce, Chengxin Zhang, Eric W. Bell, Guijun Zhang, and Yang Zhang (2022). I-TASSER-MTD: A deep-learning based platform for multi-domain protein structure and function prediction. Nature Protocols, 17. https://doi.org/10.1038/s41596-022-00728-0

Yan, Q., Lin, M., Huang, W., Teymournejad, O., Johnson, J.M., Hays, F.A., Liang, Z., Li, G., and Rikihisa, Y. (2018). *Ehrlichia* type IV secretion system effector Etf-2 binds to active RAB5 and delays endosome maturation. Proc Natl Acad Sci U S A, 115, e8977–E8986. https://doi.org/10.1073/pnas.1806904115

Yang, J., and Zhang, Y. (2015). I-TASSER server: new development for protein structure and function predictions. Nucleic Acids Res, 43, W174–181. https://doi.org/10.1093/nar/gkv342

Yu, C.S., Chen, Y.C., Lu, C.H., and Hwang, J.K. (2006). Prediction of protein subcellular localization. Proteins, 64, 643–651. https://doi.org/10.1002/prot.21018

